# Ultrasonic cerebrospinal fluid clearance improves outcomes in hemorrhagic brain injury models

**DOI:** 10.1101/2024.06.02.597001

**Authors:** Matine M. Azadian, Nicholas Macedo, Brenda J. Yu, Ryann M. Fame, Raag D. Airan

## Abstract

Impaired clearance of the byproducts of aging and neurologic disease from the brain exacerbates disease progression and severity. We have developed a noninvasive, low intensity transcranial focused ultrasound protocol that facilitates the removal of pathogenic substances from the cerebrospinal fluid (CSF) and the brain interstitium. This protocol clears neurofilament light chain (NfL) – an aging byproduct – in aged mice and clears red blood cells (RBCs) from the central nervous system in two mouse models of hemorrhagic brain injury. Cleared RBCs accumulate in the cervical lymph nodes from both the CSF and interstitial compartments, indicating clearance through meningeal lymphatics. Treating these hemorrhagic brain injury models with this ultrasound protocol reduced neuroinflammatory and neurocytotoxic profiles, improved behavioral outcomes, decreased morbidity and, importantly, increased survival. RBC clearance efficacy was blocked by mechanosensitive channel antagonism and was effective when applied in anesthetized subjects, indicating a mechanosensitive channel mediated mechanism that does not depend on sensory stimulation or a specific neural activity pattern. Notably, this protocol qualifies for an FDA non-significant risk designation given its low intensity, making it readily clinically translatable. Overall, our results demonstrate that this low-intensity transcranial focused ultrasound protocol clears hemorrhage and other harmful substances from the brain via the meningeal lymphatic system, potentially offering a novel therapeutic tool for varied neurologic disorders.

---

Impairment of cerebrospinal fluid (CSF) circulation has been shown to be both a feature of certain neurologic disorders and a pathogenic mechanism of those diseases. In the setting of both ischemic and hemorrhagic stroke, CSF and interstitial fluid clearance has been shown to be both impaired and a potential avenue for treatment. Impaired clearance of blood products in intracranial hemorrhage has been suggested as pathogenic in the setting of hemorrhagic stroke and traumatic brain injury^1–3^, with increased clearance via the meningeal lymphatics contributing to improved outcomes preclinically^4–6^. Moreover, prospective upregulation of CSF circulation whether pharmacologically or through invasive intervention yields improved outcomes both preclinically and clinically. Indeed, upregulation of meningeal lymphatic clearance through pharmacologic intervention shows improvement in certain stroke models^5,6^. Additionally, prospective CSF drainage in the setting of subarachnoid hemorrhage (SAH) improves all cause morbidity and mortality^7^ and prospective evacuation of the hematoma improves outcomes in intracerebral/intraparenchymal hemorrhage (ICH)^8^. These data suggest that a modality to drive CSF circulation could improve outcomes in varied neurologic diseases, including hemorrhagic brain injury.

Previously, we and others have shown that low intensity focused ultrasound without co-administration of exogenous agents could upregulate CSF circulation, including for driving improved brain distribution of intrathecally administered drugs^9^ and circulation of cisternally and intraparenchymally administered tracers^10,11^. While these earlier results did not confirm utility in a particular disease model, they implied that a similar low intensity ultrasound protocol could be used as a noninvasive and nonpharmacologic strategy to drive the clearance of pathogenic products that accumulate in neurologic disease and brain injury.

## Ultrasonic CSF Clearance Enhances Brain-Derived Protein and Red Blood Cell Clearance

To assess the utility of low intensity transcranial focused ultrasound (FUS) to clear soluble factors from the CSF and interstitial brain compartments, after an initial technical optimization (Figure S1) we first assessed how this FUS protocol affects young versus aged mice (Figure 1a), given that aging itself is associated both with reduced CSF bulk circulation and CSF solute clearance^12–14^. In aged mice, neurofilament light chain (NfL; a neuron-specific protein) levels in the CSF were increased relative to young adult mice consistent with previous reports^15^, and a single FUS application significantly increased NfL CSF levels compared to sham-treated controls (p ≤ 0.05; Figure 1b). This effect was not observed in young adult mice, suggesting an age-dependent effect (p > 0.05) and that the finding of FUS induced increased CSF NfL levels in aged mice was less likely to be due to direct parenchymal injury or generation of NfL, but perhaps due to mobilization of aging-related accumulation of interstitial NfL^16^. Serial FUS applications (three sessions, 24-48 hours apart) resulted in a significant decrease in CSF NfL levels in aged mice, to the level seen in young adult mice (p ≤ 0.01), with again no significant effect observed in young adults (p > 0.05) (Figure 1c), confirming that the FUS protocol does not induce neurofilament damage but rather suggests clearance of excess buildup. These results demonstrate that repeated FUS applications in this protocol can effectively clear brain-derived macromolecules from the CSF of aged mice.

**Figure 1.**
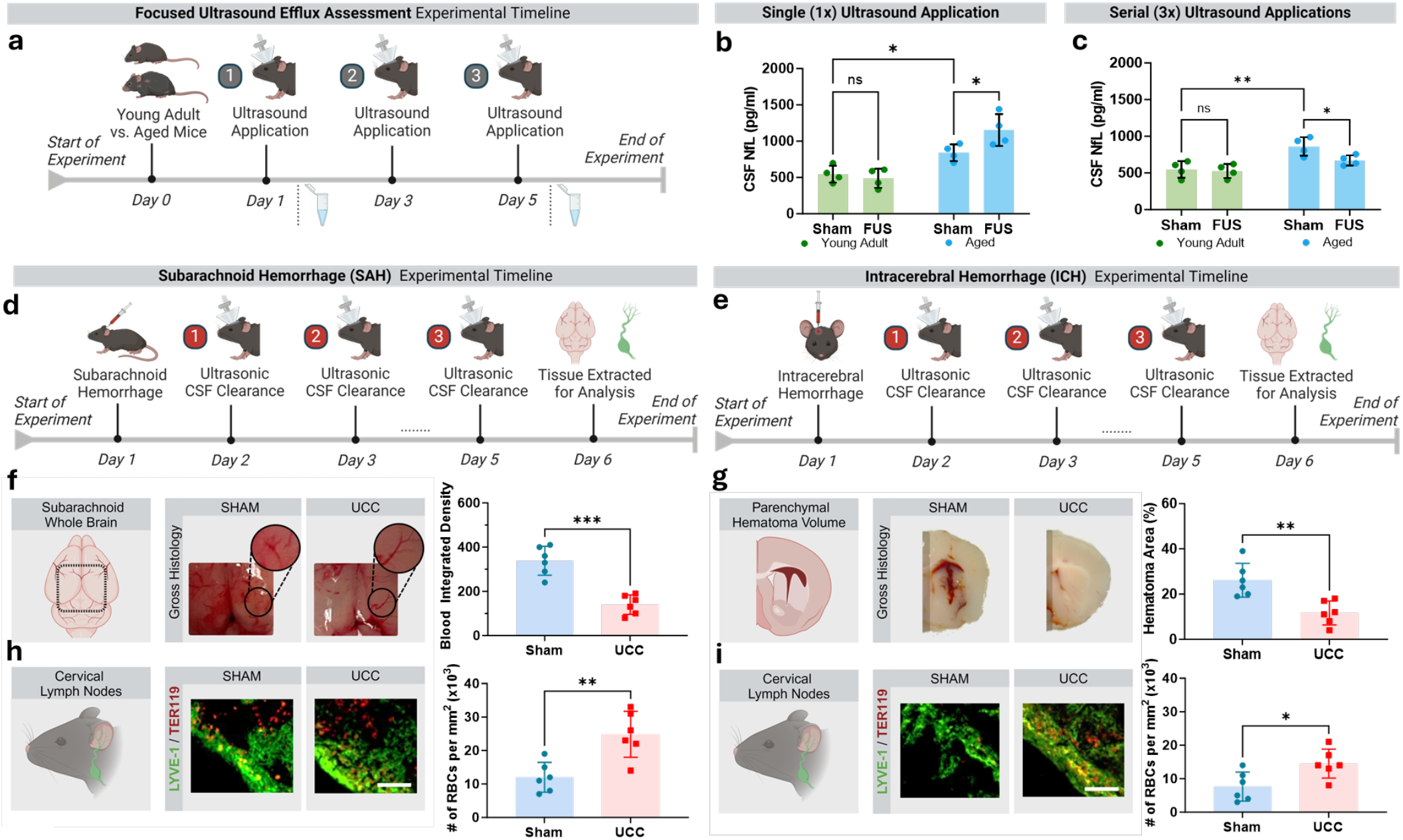
Ultrasonic CSF clearance (UCC) clears brain proteins from the CSF of aged rodents and enhances clearance of hemorrhage in the CSF and interstitial brain compartments of two hemorrhage models. **a**. Timeline of focused ultrasound (FUS; 250k Hz, 0.45 MPa, 20% duty-cycle, 50 ms pulse width) efflux experiment. **b**. A single FUS application increases neurofilament light chain (NfL) in the CSF of aged mice compared to sham, with no effect on young adult mice. **c**. Serial FUS application (3x, 24-48 hours apart) decreases neurofilament light chain (NfL) in the CSF of aged mice compared to sham, with no effect on young adult mice. **d**. Timeline of subarachnoid hemorrhage (SAH) experiment. **e**. UCC clears CSF dispersed red blood cells (RBCs) compared to sham. **f**. Greater accumulation of RBCs in the cervical lymph nodes of SAH mice with UCC compared to sham. **g**. Timeline of intracerebral hemorrhage (ICH) experiment. **h**. UCC clears interstitial RBCs compared to sham. **i**. Greater accumulation of RBCs in the cervical lymph nodes of ICH mice with UCC compared to sham. Data presented as mean ± S.D; b-c: two-way ANOVA with post-hoc uncorrected Fisher’s LSD, f-i: unpaired (two-tailed) t-tests with Welch’s correction. *: p ≤ 0.05, **: p ≤ 0.01, ***: p ≤ 0.001. n = 4 for each group.

Given these promising results, we next looked to determine the impact of this ultrasonic CSF clearance (UCC) protocol in models of neurologic disease. We specifically turned our attention to hemorrhagic brain injury as impaired clearance of hemorrhagic products is thought to be pathogenic in both subarachnoid hemorrhage (SAH) and intracerebral (also known as intraparenchymal) hemorrhage (ICH)^17,18^. In a mouse SAH model, UCC treatment significantly reduced the amount of red blood cells detected in the CSF compared to sham treatment (p ≤ 0.001) (Figure 1e). This reduction was associated with a greater accumulation of red blood cells in the cervical lymph nodes, indicating efficient clearance through the meningeal lymphatic system (p ≤ 0.01) (Figure 1f). Similarly, in the ICH model, UCC treatment led to a significant reduction in interstitial red blood cells compared to sham (p ≤ 0.01) (Figure 1h). This clearance was again accompanied by increased red blood cell accumulation in the cervical lymph nodes (p ≤ 0.05) (Figure 1i). These findings suggest that UCC facilitates the clearance of hemorrhagic debris from both the CSF and interstitial compartments, likely through the meningeal lymphatic pathway.

## UCC Reduces Post-Hemorrhagic Neuroinflammatory and Neurocytotoxic Profiles

Neuroinflammation and neurocytotoxicity are critical factors that contribute to poor outcomes following hemorrhagic brain injury. To evaluate the impact of UCC on these processes, we performed immunohistochemical analyses of brain tissue from UCC-treated SAH and ICH models. UCC treatment significantly reduced microglial activation, as indicated by lower IBA-1 staining in both models (p ≤ 0.05) (Figure 2a-c). Additionally, astrocytic activation, assessed by GFAP staining, was also significantly reduced in UCC-treated animals in both models (p ≤ 0.05) (Figure 2c). These results indicate that UCC effectively attenuates neuroinflammatory responses following hemorrhagic brain injury. Furthermore, histological assessment of neuronal degeneration using Fluoro-Jade C (FJ-C) staining revealed a significant reduction in degenerating neurons in UCC-treated animals compared to sham (p ≤ 0.01) (Figure 2d). This reduction in neurotoxicity highlights the protective effects of noninvasive early blood clearance post-hemorrhage in reducing neuronal damage.

**Figure 2.**
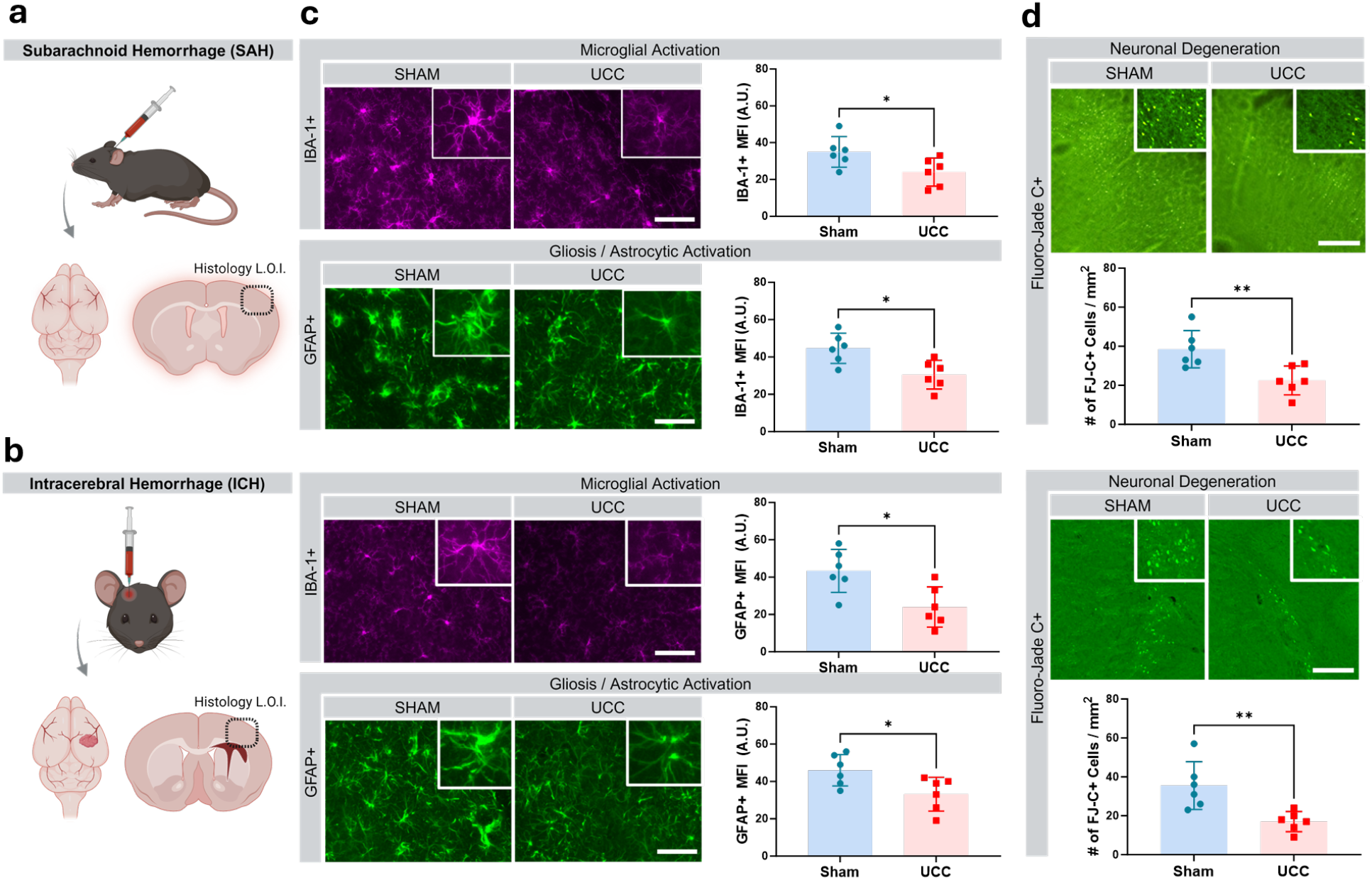
UCC reduces neuroinflammation and neurocytotoxicity post-hemorrhagic brain injury. **a**. Schematic of subarachnoid hemorrhage (SAH) model with the location of interest (LOI) sampled for histological analyses. **b**. Schematic of the intracerebral hemorrhage (ICH) model with the LOI sampled for histological analyses. **c**. Immunohistological assessment of both SAH and ICH brains for microglial activation (IBA-1, purple) and gliosis/astrocytic activation (GFAP, green) revealed a decrease in both neuroinflammatory markers in both models of hemorrhagic brain injury with UCC compared to sham. **d**. Histological assessment of both SAH and ICH brains for neuronal degeneration (FJ-C, green) revealed a decrease with UCC in both models compared to sham. Data presented as mean ± S.D; unpaired (two-tailed) t-tests with Welch’s correction. *: p ≤ 0.05, **: p ≤ 0.01. n = 6 for each group.

## Improved Behavioral Outcomes and Survival with UCC Treatment

Next, we assessed functional recovery and survival as indicators of the efficacy of UCC as a potential therapeutic intervention for brain injury. Our pilot studies indicated that the SAH model did not produce a sustained behavioral deficit. Therefore, to assess the impact of UCC on behavioral outcomes, we conducted the corner turn test and forelimb grip strength assessments using the ICH model (Figure 3a). UCC treatment significantly improved both functional outcomes as early as six days post-hemorrhage, with sustained improvements observed up to 14 days (p ≤ 0.01) (Figure 3b).

**Figure 3.**
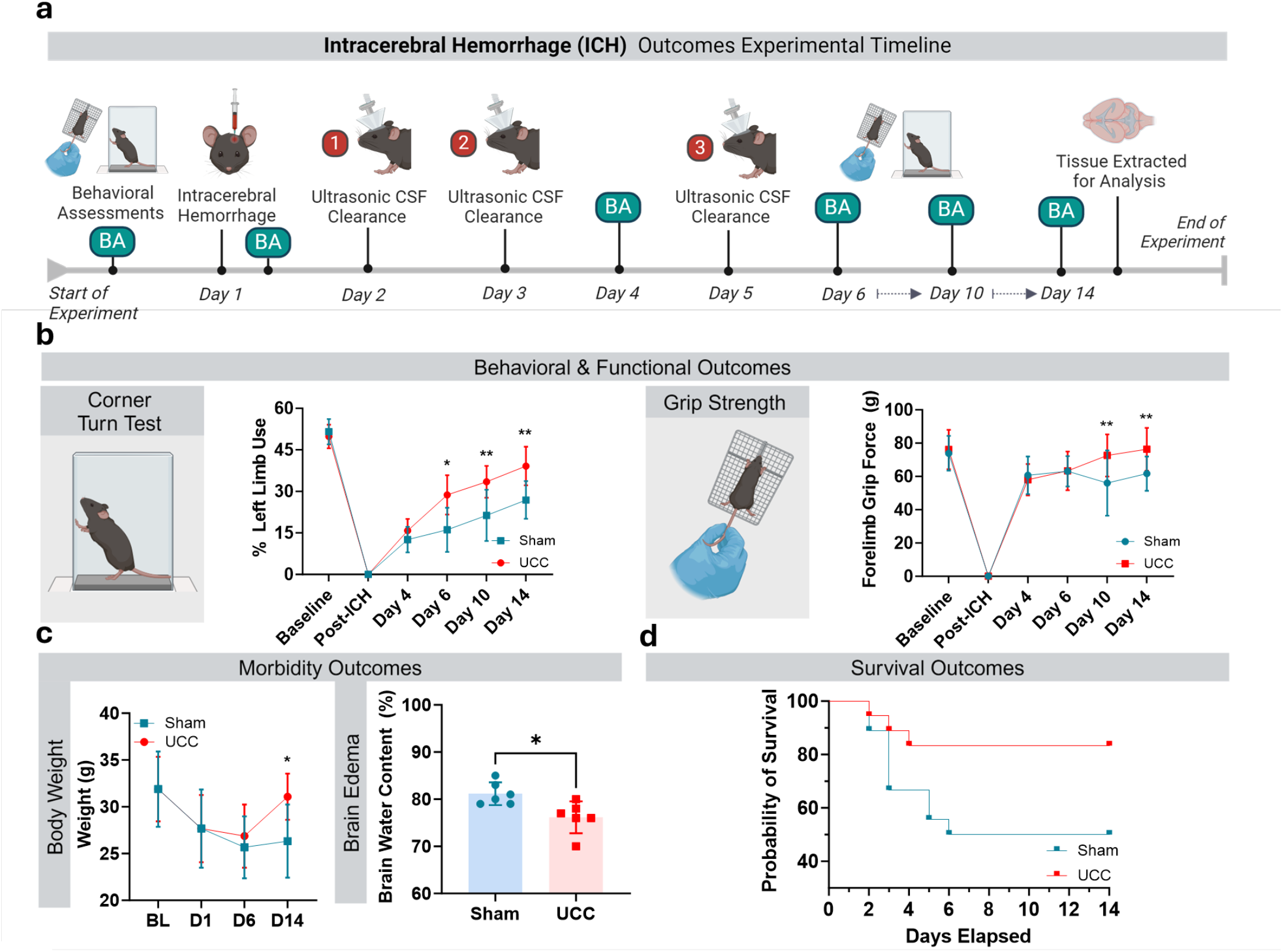
UCC improves behavioral outcomes and reduces morbidity and mortality post-hemorrhagic brain injury a. Timeline of intracerebral hemorrhage (ICH) outcomes experiment. **b**. Behavioral/functional assessments (corner turn test and grip strength) revealed an improvement in functional outcomes as early as 6 days post-hemorrhage with UCC compared to sham. **c**. Decreased morbidity outcomes (body weight and brain water content, edema) with UCC compared to sham. **d**. Increased survival with UCC intervention compared to sham, χ^2^(2) = 4.08, p < 0.05. Data presented as mean ± S.D.; b-c: mixed-effects model (REML) with post-hoc Sidak’s multiple comparisons test; c, left: two-way ANOVA with post-hoc Sidak’s multiple comparisons test; c, right: unpaired (two-tailed) t-tests with Welch’s correction; d: log-rank (Mantel-Cox) test. *: p ≤ 0.05, **: p ≤ 0.01. n=18 for each group; n=6 for brain water content experiment.

Morbidity outcomes, including body weight and brain water content (indicative of edema), were also assessed. UCC-treated mice exhibited significantly lower brain water content and recovered their body weight faster compared to sham-treated controls (p ≤ 0.05) (Figure 3c). Notably, UCC treatment significantly increased survival rates compared to sham (p < 0.05) (Figure 3d). These disparate survival rates between the sham and UCC groups underscore the positive behavioral effects (Figures 3a-b), which were observed despite a survivorship bias in these data that otherwise serves to lessen the observable behavioral effect sizes. These findings demonstrate that UCC not only enhances functional recovery but also reduces morbidity and mortality following hemorrhagic brain injury.

## UCC Acts Through Molecular Mechanotransduction Pathways

To elucidate the underlying mechanisms of UCC-mediated clearance, we investigated the role of mechanosensitive ion channels in the ability of UCC to exert its protective effects (Figure 4a). At the intensity, duty cycle, and ultrasound frequency used here, it is unlikely that significant (>0.5 °C) parenchymal heating is being induced in these experiments. Accordingly, we hypothesized that ultrasound is acting mechanically to clear the CSF in this protocol. Possible mechanisms for this mechanical effect are direct induction of a convection or oscillation of the CSF and interstitial fluid with or without an accompanying mechanical change of the matrix to then drive downstream CSF circulation^19^. However, our prior results demonstrated that the response to this protocol extends for a substantial amount of time beyond the direct ultrasound exposure^9^, implying induction of a prolonged cascade of events leading to longer-term CSF circulation upregulation. Therefore, we instead hypothesized that actuation of mechanosensitive channels could contribute to increased CSF clearance as these channels play critical roles in cellular responses to mechanical stimuli like focused ultrasound^20,21^. Additionally, agonism of these channels yields meningeal lymphatic vessel sprouting with resultant increased CSF egress^22,23^. To determine the relative contribution of these mechanosensitive channels to the effect of UCC, we cisternally administered a mechanosensitive ion channel blocker (GsMTx4, a selective blocker of Piezo and TRP channels) in the ICH model to determine its effect on UCC-mediated clearance (Figure 4a). Blockade of mechanosensitive channels with GsMTx4 significantly reduced the efficacy of blood product clearance by UCC, as evidenced by increased hematoma volumes in the UCC+GsMTx4 treatment group versus UCC treatment alone (p ≤ 0.01) (Figure 4b). This finding indicates that mechanosensitive ion channels mediate the majority of the observed clearance effects of UCC.

**Figure 4.**
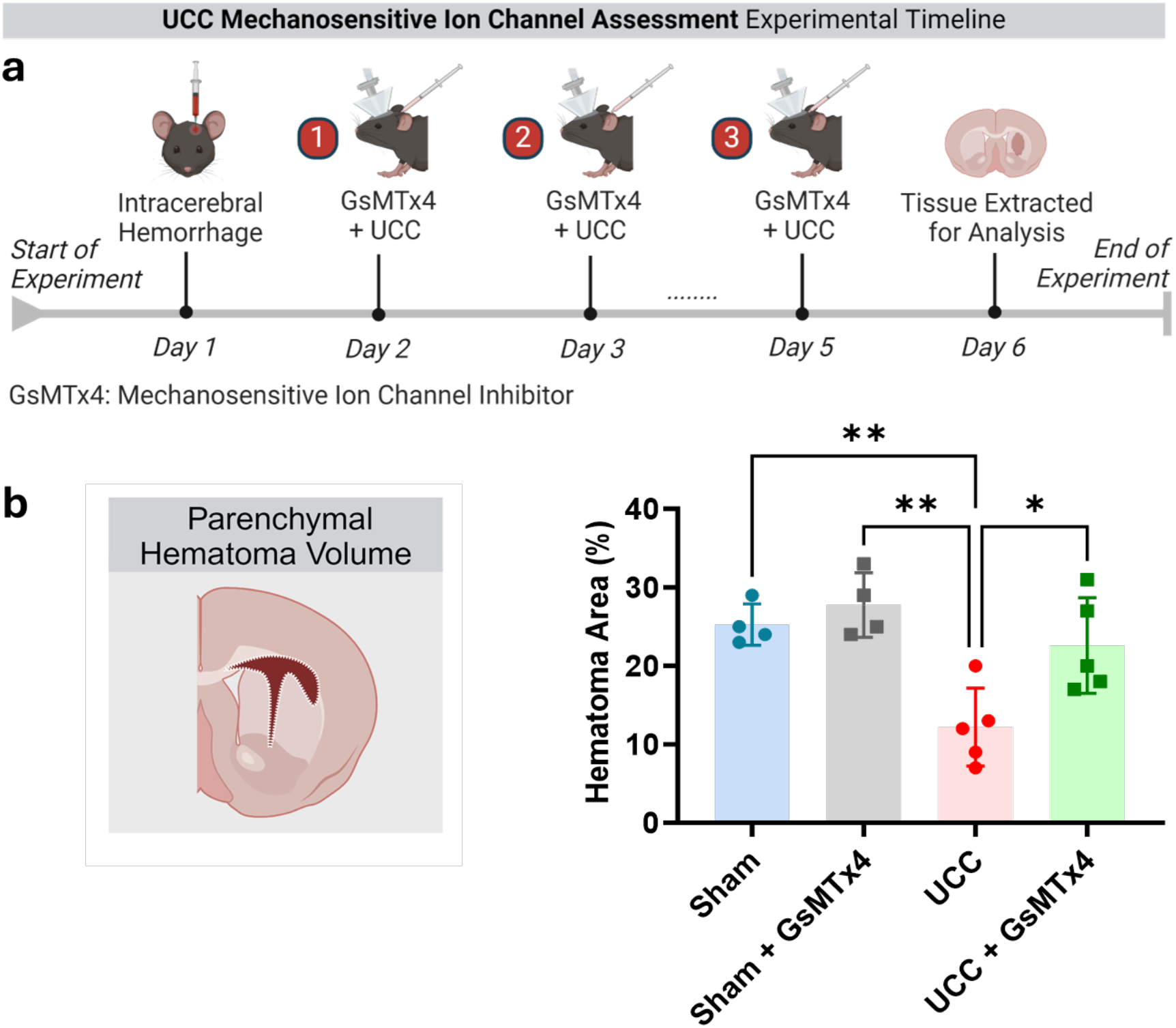
UCC is reversed by a mechanosensitive ion channel blocker. **a**. Timeline of intracerebral hemorrhage (ICH) experiment with UCC and a mechanosensitive ion channel blocker (GsMTx4). **b**. Administration of a GsMTx4 reduced the blood product clearance of UCC, as assessed by hematoma volumes across treatment groups. Data presented as mean ± S.D.; one-way ANOVA with post-hoc Tukey’s multiple comparisons test. *: p ≤ 0.05, **: p ≤ 0.01. n = 4 mice for each group.

Overall, our results demonstrate that UCC indeed facilitates the clearance of pathogenic substances from the brain, thereby reducing neuroinflammation and neurocytotoxicity, and improving functional outcomes and survival following hemorrhagic brain injury. We have experimentally confirmed that UCC acts via mechanosensitive channel transduction to mediate these effects, indicating that UCC acts as a noninvasive and nonpharmacologic agonist of molecular mechanotransduction pathways to elicit increased CSF circulation and egress.

Notably, the design of the UCC protocol enables a ready path towards clinical translation. The 250 kHz ultrasound frequency used here is in the range of commercial clinical transcranial FUS systems currently on market worldwide^24^ and is further enabled by having relatively low attenuation and aberration when transmitting this ultrasound frequency through the human skull^25^. Additionally, the estimated in situ peak negative pressure used in these experiments corresponds to a mechanical index (MI) of 0.9, which is well within FDA guidelines for safe application^26^, indicating that a clinical application of UCC should be able to receive a nonsignificant risk (NSR) designation, further enabling clinical translation. In addition, UCC does not require co-administration of exogenous agents like nanoparticles or microbubbles, further uncomplicating its eventual application. Finally, as its mechanism does not require a particular pattern of brain activity (i.e., it is effective in anesthetized animals), it should be applicable in humans in a range of cognitive states including while awake, asleep, engaged in some other task, or obtunded in the immediate post-injury recovery period.

Furthermore, UCC provides a method to drive CSF circulation and egress through a complementary mechanism to pharmacologic approaches and approaches that activate particular neural activity patterns that are correlated to this circulation^27,28^. Accordingly, UCC provides an alternate method for understanding the role of CSF circulation and egress in the variety of disorders it has been implicated including sleep^29^, neurodegeneration^30^, traumatic brain injury^31^, and mental health^32^, that is non-pharmacologic and could be applied to subjects in varied states of cognition or wakefulness. Indeed, further validation of UCC and similar approaches for harnessing CSF clearance in the variety of disorders to which it has been implicated could and should be applied in appropriate clinical trials.

## Methods

### Animals

All experiments were approved by the Institutional Animal Care and Use Committees of Stanford University. Experiments were conducted with adult (3-4 month) and aged (18-20 month) male C57BL/6NCrl mice with bodyweight 30-40 g (Charles River Laboratories, Wilmington, MA, USA). All mice were housed in a temperature-controlled (22 °C), humidity-controlled (33–39%) environment under a 12h /12 h light-dark cycle and were provided with food and water ad libitum. Mice were randomly assigned to one of two treatment groups: (1) no treatment (sham), and (2) treatment with a focused ultrasound protocol, herein referred to as ultrasonic CSF clearance (UCC).

### Ultrasonic CSF Clearance Protocol

Focused ultrasound (250kHz, 0.45MPa, 50ms pulse-width, 20% duty cycle for 10 min; UCC) or sham (ultrasound power off) was applied transcranially throughout the brain (Fig. S2). Before application, the fur on the head was removed using chemical hair depilatory. The transducer was coupled with ultrasound gel to the dorsal surface of the head, and ultrasound applied while the mice were anesthetized under ketamine/xylazine (90 mg/kg and 10 mg/kg, respectively). Body temperature, cardiac and respiratory rates, and O2 saturation were monitored throughout the experiment. Environmental heating was used to help maintain body temperature.

### Hemorrhagic Models

Two previously characterized models of hemorrhagic stroke, subarachnoid (SAH)^33^ and intracerebral (ICH)^34^, were implemented in this study. Briefly, for the SAH model, 25µl of autologous blood was withdrawn from the tail vasculature into heparinized capillary tubing and injected into the cisterna magna. For the ICH model, 25 µl of autologous blood was withdrawn from the tail vasculature and injected into the right striatum. Mice were anesthetized under ketamine/xylazine (90 mg/kg and 10 mg/kg, respectively). Body temperature, cardiac and respiratory rates, and O2 saturation were monitored throughout the experiment. Environmental heating was used to help maintain body temperature.

### Cerebrospinal Fluid Collection

Cerebrospinal fluid was collected by inserting a pulled glass capillary, connected to an aspirator tube, into the cisterna magna. Collected CSF was centrifuged at 1,000 x g for 10 min at 4°C to remove any cellular debris. Neurofilament light polypeptide (NEFL) levels were quantified by a high-sensitive ELISA (Cat# EKU10307, Biomatik, Ontario, Canada). Tests were performed in duplicate using 10 uL CSF diluted in aCSF (Tocris Biosciences, Minneapolis, MN, USA). Absorbances were quantified using a Spark Multimode Microplate Reader (Tecan, Männedorf Switzerland).

### Histology and Immunostaining

Mice were euthanized and fixed via transcardial perfusion with phosphate-buffered saline (PBS) and 4% paraformaldehyde (PFA) at timepoints described in each experimental timeline (Fig 1-4). Brains were fixed in 4% PFA for 24 h, cryoprotected by serial incubation in 15% and 30% sucrose solutions for 24 h each, and then frozen at −80 °C in optimal cutting medium (OCT) compound. Brains were sectioned at 30 μm thickness using a cryostat (LEICA CM 1950, Buffalo Grove, IL, USA). Every 5^th^ section (150 μm apart) was collected for imaging. The specimen temperature was set at −21 °C. Tissue sections were stored in cryoprotectant solution prior to staining. For immunofluorescence, the free-floating tissue sections were blocked by 0.3% PBST with 10% normal goat serum for 1 h at room temperature, then incubated with primary antibodies overnight at 4 °C. After washing with PBS three times for 5 min each, sections were incubated in secondary antibodies for 2 h at room temperature. The primary antibodies used in immunofluorescence included rabbit anti-LYVE-1 antibody (1:500; Abcam, Cat. No. ab14917), rat anti-TER-119 antibody (1:500; Invitrogen, Cat No. 14-5921-82), recombinant rabbit anti-IBA-1 (1:500; Abcam, Cat No. ab178846), and chicken anti-GFAP antibody (1:500; Abcam, Cat. No. ab4674). The corresponding secondary antibodies were used as follows: AlexaFluor 488-labeled goat anti-rabbit antibody (1:500; ThermoFisher Scientific, Cat. No. A32731), AlexaFluor 555-labeled goat antirat antibody (1:500; ThermoFisher Scientific, Cat. No. A48263), AlexaFluor 647-labeled goat anti-rabbit antibody (1:500; ThermoFisher Scientific, Cat. No. A32733), and AlexaFluor 488-labeled goat anti-chicken antibody (1:500; ThermoFisher Scientific, Cat. No. A32931). For neurodegeneration staining, tissue sections were incubated in 0.06 % potassium permanganate for 20 min and in 0.0001 % Fluoro-Jade C (Biosensis, Thebarton, SA, Australia). Tissue sections were mounted on microscope glass slides (Fisher, Pittsburgh, PA) and cover-slipped with ProLong Gold Antifade Mountant (Thermo Fisher Scientific, Waltham, MA, USA). All images were collected with a fluorescence microscope (BZ-X800, Keyence Corp., Itasca, IL, USA) and processed with BZ-X Advanced Analysis Software (Keyence Corp., Itasca, IL, USA) or ImageJ Software (Version 1.53, National Institutes of Health, Bethesda, MD, USA). For hematoma volume, the mean area of blood was calculated in coronal segments using ImageJ and reported as percent hematoma area over total hemispheric area. For immunofluorescence quantification, signals above manually thresholded background were used for ROI segmentation to calculate total mean fluorescence intensity using BZ-X Advanced Analysis Software. All imaging and thresholding parameters were completed by personnel blinded to experimental conditions.

### Neurobehavioral Function Evaluation

Two behavioral tests were used to evaluate behavioral and functional outcomes as previously described: a corner turn test and a grip strength test^35^. Briefly, for the corner turn test, quantification of turning preference upon approaching a 30° corner was used to assay sensorimotor deficits. The values were calculated as percent of left versus right limb use on turn for a total 3 min. duration per session. For the grip strength test, motor function was assessed via the peak force (G) required for mice to release their grip from a grid bar as quantified by a digital grip strength meter (Maze Engineers, Skokie, IL, USA). The average of 3 attempts was calculated per test session. All behavioral assessments were completed by personnel blinded to experimental conditions.

### Morbidity Evaluation

Total body weight (g) and brain water content (edema) percentage were used as indicators of post-hemorrhagic morbidity. Brain water content was assessed at day 14 post-hemorrhage. The wet weight of each brain was recorded following euthanasia and extraction. The brains were then dehydrated at 110°C for 72h. Dry weights were recorded, and brain edema was evaluated as the difference in percent brain water content.

### Reagents

For mechanosensitive ion channel inhibition, we mixed GsMTx4 (MedChem Express, Cat. No. HY-P1410) with artificial CSF and injected into the cisterna magna of mice 0.5 h prior to focused ultrasound application.

## Acknowledgments

We would like to thank Paul M. George and Jeremy J. Heit for a variety of useful discussions and the use of key reagents and equipment. We would like to thank Ronald Watkins and the whole Airan Lab for helpful discussions.

## Author contributions

Conceptualization: RDA

Methodology: MMA, NM, BJY, RMF, RDA

Investigation: MMA, RDA

Visualization: MMA, RDA

Funding acquisition: RDA

Project administration: RDA

Supervision: RDA

Writing – original draft: MMA, RDA

Writing – review & editing: MMA, RMF, RDA

## Data and materials availability

Raw data underlying all figures are available via a source data file submitted with this manuscript. Any additional data not presented is available from the corresponding author upon reasonable request.

