## Supplementary Figure S1 for "Ultrasonic cerebrospinal fluid clearance improves outcomes in hemorrhagic brain injury models"

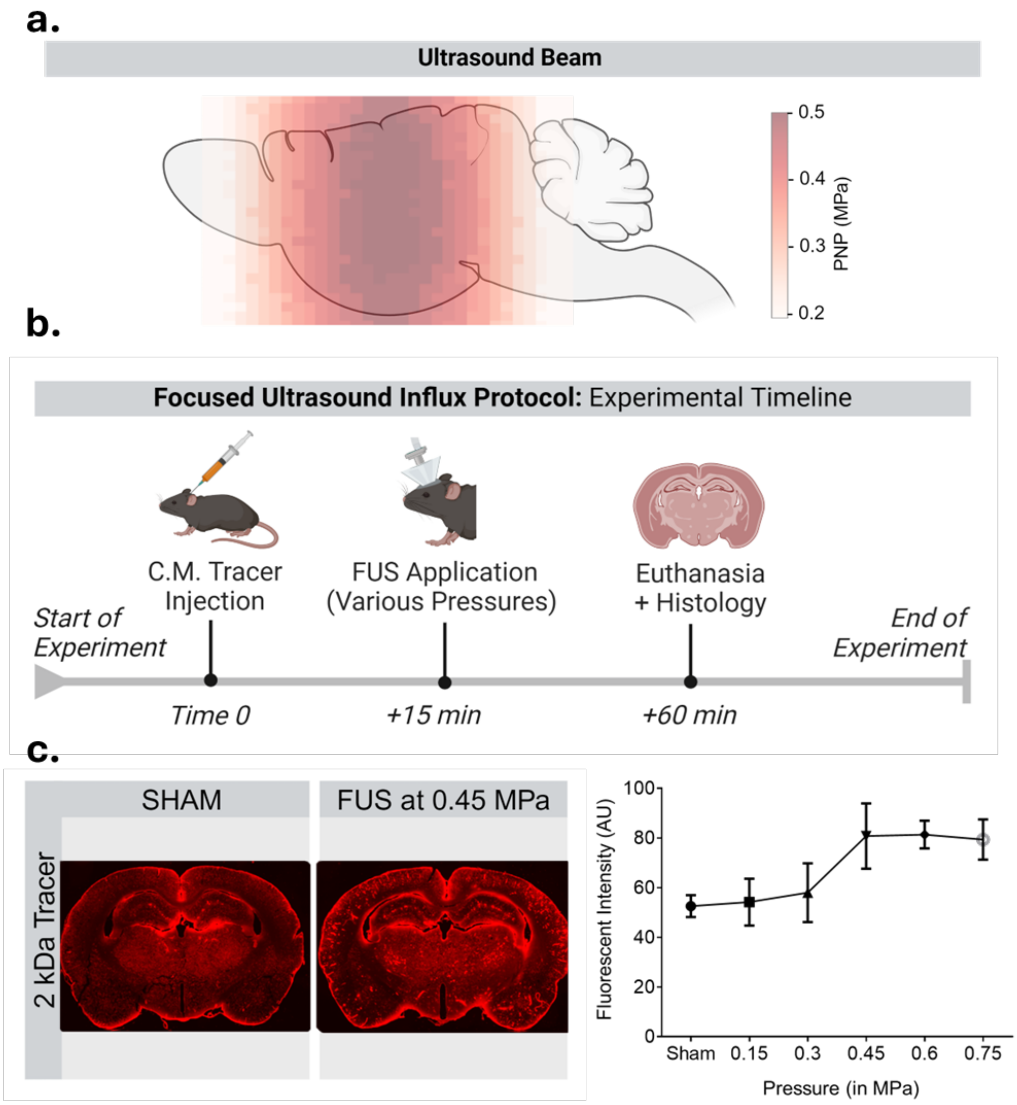


**Figure S1. Focused ultrasound induced influx effects peak at 0.45 MPa peak negative pressure for 250 kHz ultrsasound**

**a.** Schematic of ultrasound beam and estimated peak negative pressure (PNP) intensities across a sagittal outline of the mouse brain. **b.** Timeline of PNP parameter assessment experiment. **c.** Representative coronal brain slice and quantification of mean fluorescent intensity following intracisternal injection of a 2kDa tracer dye at 60 min post-injection. Given the inflection at 0.45 MPa estimated in situ peak negative pressure with saturation of the effect at higher pressures, this pressure was chosen for subsequent analysis. Data presented as mean ± S.D.; n = 3 mice at each pressure measure.
